# Influence of season, tourist activities and camp management on body condition, testicular and adrenal steroids, lipid profiles, and metabolic status in captive bull elephants in Thailand

**DOI:** 10.1101/507855

**Authors:** Treepradab Norkaew, Janine L. Brown, Pakkanut Bansiddhi, Chaleamchat Somgird, Chatchote Thitaram, Veerasak Punyapornwithaya, Khanittha Punturee, Preeyanat Vongchan, Nopphamas Somboon, Jaruwan Khonmee

## Abstract

We previously found relationships between body condition and physiological function affecting health and welfare of female tourist camp elephants in Thailand, and used that approach to conduct a similar study of bulls elephants in the same camps (n = 13). Elephants received a body condition score (BCS) every other month, and fecal glucocorticoid metabolite (FGM) concentrations were measured in twice monthly samples for 1 year. Effects of season, camp management and tourist activity on lipid profiles [total cholesterol (TC), low density lipoproteins (LDL), high density lipoproteins (HDL), triglycerides (TG)] and metabolic factors [insulin, glucose, fructosamine, glucose to insulin ratio (G:I)] were determined and correlated to measures of body condition, testosterone and adrenal function. Positive correlations were found between BCS and TG, between FGM and TG, HDL and glucose, and between testosterone and HDL (p<0.05), whereas BCS and testosterone were both negatively associated with the G:I (p<0.05). Elevated FGM concentrations were associated with altered lipid profiles and metabolic status and were higher in winter compared to summer and rainy seasons. Insulin and glucose levels were higher, while the G:I was lowest, in the winter season. Strong positive associations (p<0.001) were found between TC and HDL, LDL and HDL and glucose, and glucose and insulin. By contrast, negative relationships (p<0.001) were found between the G:I and HDL and glucose, and between insulin and G:I. Differences also were found between High and Low tourist season months for FGM, insulin, and G:I, which may be related to tourist activities. Last, there was notable variation among the camps in measured parameters, which together with the tourist season effect suggests camp management related to tourism may affect physiological function and welfare; some negatively like feeding high calorie treats, but others are positively, like exercise. Last, compared to females, bulls elephants appear to be in better physical health.

## Introduction

The Asian elephant (*Elephas maximus*) has been listed as endangered in Appendix 1 of the Convention on International Trade in Endangered Species of Wild Fauna and Flora (CITES 2018) since 1973, with wild populations declining in several range countries. Some captive elephant populations in Asia also are not sustaining, due in part to low reproductive and high mortality rates. As elephants are a long-lived species that produce only a few calves in their lifetime, it is important to better understand factors affecting health and reproduction to prevent further population declines. Considerable information now is available about the basic biology of elephants, especially the reproductive physiology of females, while bulls have received comparatively less attention (e.g., [1]).

In recent years, studies have focused on associations between health and reproduction in captive elephants, particularly females, with problems linked at least in part to obesity because of too little exercise, diets that are too high in calories, or both [2–4]. There are numerous documented links between obesity, metabolic and lipid problems in other species, including humans, companion and domestic animals [5–7]. Abnormally high blood glucose, triglycerides (TG), and cholesterol have been linked to a number of health issues, collectively called metabolic syndrome, such as hypertension, hyperlipidemia, insulin resistance, and type 2 diabetes [8, 9]. In zoo Asian and African elephant females, negative associations have been found between body condition scores (BCSs) and the glucose to insulin ratio (G:I) [4], suggesting that, as is the case in women, low G:I reflects an unhealthy state [10]. Recently, Norkaew *et al*. [11] found high BCSs also were associated with a low G:I in captive female Asian elephants in Thailand, as well as higher total cholesterol (TC) and low density lipoprotein (LDL). To our knowledge, only one group has measured glucose and insulin in bull Asian elephants, and found a significant sex difference; mean G:I was 143 units lower in females than males, which corresponded to higher BCSs [12]. Taken together, evidence suggest that being overweight may lead to potentially negative health consequences, at least in females, leading to questions about whether bull elephants are prone to similar obesity-related risks.

In addition to body condition, stress can affect metabolic health and lipid parameters. Glucocorticoids (GCs) are endogenous stress hormones that affect nearly every organ and tissue in the body, modulating various physiological processes including energy homeostasis (metabolism), immunology, behavior, reproductive function, cell proliferation, and survival [13]. While important for maintaining metabolic equilibrium, excessive GC exposure for prolonged periods can have devastating effects on health [14]. Working elephants in Thailand interact with the public in a variety of ways, including performing in shows, trekking, bathing, and painting. Often these activities are not closely monitored or regulated, and could be sources of stress to individual animals. Tourist activities have been shown to compromise welfare and negatively affect behavior and physiology in other species, resulting in increased hiding behaviors [15, 16], heightened vigilance [17, 18], stereotypies [19, 20], poor body condition [21], and elevated cortisol [22, 23]. In a recent study, stress levels based on fecal GC metabolite (FGM) measures, and several lipid and metabolic factors were higher in females elephants during the high tourist season, suggesting some tourist activities may have a negative impact on health and well-being (Norkaew *et al*., [24]. However, there appeared to be a protective effect of exercise, with hours of daily exercise being associated with better body condition and lipid and metabolic profiles, so elephant trekking per se may not be as bad for welfare as some animal activist organizations claim [25].

Therefore, the aims of this study were to examine: 1) relationships between BCS and FGM on metabolic function (insulin, glucose, fructosamine, G:I) and lipid profiles (TC, TG, HDL, LDL) in male Asian elephants in Thailand; 2) the effect of age on FGM, metabolic function and lipid profiles; and 3) how camp management and the tourist season affects adrenal, lipid and metabolic function in working bull elephants. In addition, because obesity has been shown to be related to declining circulating testosterone levels [26], and reductions in sex hormone binding globulin (SHBG) associated with hyperinsulinemia in obese men [27], we also examined lipid and metabolic relationships with serum testosterone levels. Understanding how management affects health could aid in developing science-based strategies to create sustainable populations of elephants that take into consideration both physical and psychological welfare needs.

## Materials and methods

### Environmental data

Weather in Thailand is hot and humid, with three official seasons: summer (16 February–15 May), rainy (16 May–15 October) and winter (16 October–15 February). Information on daily temperature (°C), amount of rainfall (mm/day), and humidity (%), averaged by month, was obtained from The Northern Meteorological Center, Meteorological Department, Ministry of Information and Communication Technology, Chiang Mai, Thailand [28]. A thermal-humidity index (THI) was calculated based on air temperature and relative humidity using the following formula: THI = (1.8×*T*db+32) − (0.55–0.0055×RH) × (1.8×Tdb–26), where the Tdb is the temperature of air measured by a thermometer freely exposed to the air, but shielded from radiation and moisture, and RH is the relative humidity (%) [29]. High (November – February) and Low (March – October) tourist seasons were defined by the Tourism Authority of Thailand.

### Animals

This study was approved by the Faculty of Veterinary Medicine, Chiang Mai University, Animal Care and Use Committee (FVM–ACUC; permit number S39/2559). Table 1 describes the elephants and tourist camp activities in this study, which are a subset of camps described in Norkeaw *et al*. [11]. Thirteen adult male Asian elephants (age range, 16–50; mean, 35.1 ± 2.9 years) were housed at three tourist camps within 43–72 km of the Faculty of Veterinary Medicine, Chiang Mai University (latitude 18°47’N, longitude 98°59’E, altitude 330 m). Tourists interacted with elephants through riding programs (bareback or with a saddle) and feeding of supplementary foods. Elephants were fed primarily corn stalk, *napier grass* (*Pennisetum purpureum*) and bana grass (*Pennisetum purpureum* X, *P. americanum* hybrid) with unlimited access to fresh water. Animals were given an annual physical examination by staff veterinarians, and were in good health during the study.

**Table 1.**
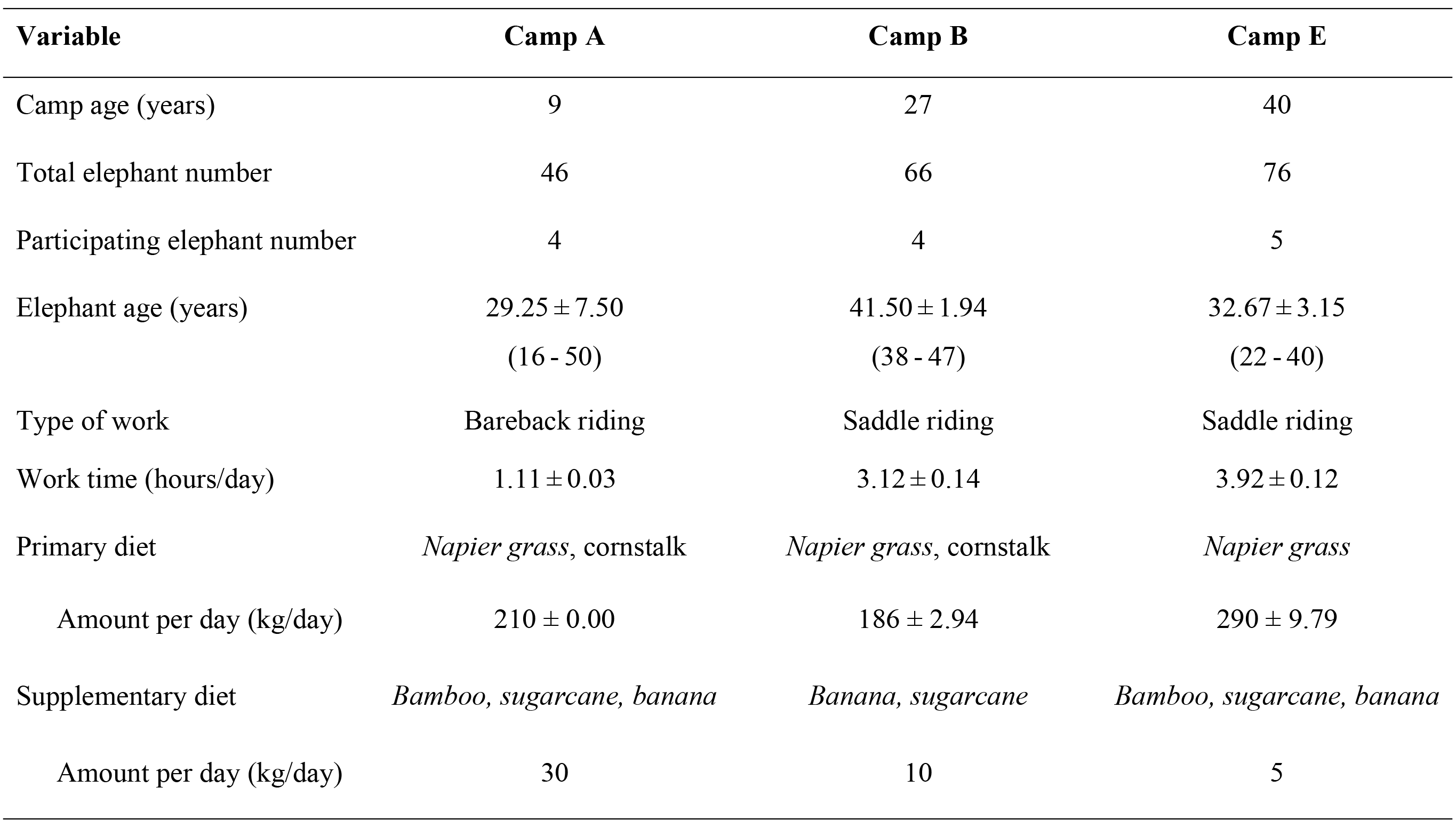
Description of elephant camps in the study. Information includes number of years the camp has been in operation (camp age), total number of elephants in each camp, number of elephants participating in the study, participating elephant mean (± SEM) and range age and work time, the general type of work conducted with tourists, and estimated amounts of primary and supplemental food items provided.

### Body condition scoring

Once every 2 months, rear and side view photographs were taken of each elephant to permit a visual evaluation of the backbone, rib bone and pelvic bone areas, and scored 1–5 (1 = thinnest; 5 = fattest) in 0.5-point increments as described by Norkaew *et al*. [11]. All photos were evaluated by three experienced elephant veterinarians, and the scores averaged. Intra-class correlations determined the inter-assessor reliability was 0.85.

### Blood collection

Blood samples (10 ml) were collected from each elephant from an ear vein by elephant camp staff or Chiang Mai University veterinarians twice monthly for 1 year. All elephants were conditioned to the blood sampling procedure; however, for safety reasons, blood collection was not attempted on any bulls exhibiting signs of musth. Three bulls came into musth during October and December, for 2 – 3 months each. Blood was centrifuged at 1,500 x g for 10 minutes within a few hours of collection, and the serum stored at -20°C until processing and analysis.

### Metabolic and lipid marker analysis

Metabolic and lipid markers were analyzed as described by Norkaew *et al*. [11]. Serum glucose was measured using an automated glucose analyzer (Glucinet T01-149, Bayer, Barcelona, Spain), serum fructosamine was measured by a colorimetric method in a Biosystems BA400 clinical chemistry analyzer (©Biosystems S.A., Barcelona, Spain), and a validated solid-phase, two-site bovine insulin enzyme immunoassay (EIA; Cat. No. 10-1113-01; Mercodia, Uppsala, Sweden) was used to measure serum insulin concentrations. All samples were analyzed in duplicate; intra- and inter-assay CVs were <10% and <15%, respectively.

Serum lipids were quantified using a Mindray BS Series analyzer (Mindray BS-380, Shenzhen Mindray Bio-Medical Electronics Co., Ltd.), total cholesterol was measured by a cholesterol oxidase-peroxidase (CHOD-POD) method, and triglycerides were measured by a glycerokinase peroxidase-peroxidase (GPO-POD) method.

### Fecal extraction and GC metabolite analysis

Fecal samples were collected immediately after defecation, and about 50 g of well-mixed sub aliquots were placed in plastic ziplock bags, and stored on ice until transported to freezers at the Chiang Mai University. The fecal extraction technique is described in Norkaew *et al*. [11]. Briefly, samples were dried in a conventional oven at 60°C for ~24-48 hours and stored at -20°C until extraction. Frozen dried fecal samples were thawed at room temperature (RT), mixed well and 0.1 g (± 0.01) of dry powdered feces extracted twice in 90% ethanol in distilled water by boiling in a water bath (96°C) for 20 minutes and adding 100% ethanol as needed to keep from boiling dry. Samples were centrifuged at 1,500 x g for 20 min, and the combined supernatants dried under air in a 50°C water bath. Dried extracts were reconstituted in methanol, diluted 1:3 in assay buffer (Cat. No. X065, Arbor Assays, Arbor, MI, USA) and stored at -20°C until enzyme immunoassay (EIA) analysis.

Concentrations of FGM were determined using a double-antibody EIA with a polyclonal rabbit anti-corticosterone antibody (CJM006) validated for Asian elephants [30] and described by Norkaew *et al*. [11]. Samples (50 μl) or corticosterone standards (50 μl) were added to appropriate wells. Corticosterone-horseradish peroxidase (HRP) (25 μl; 1:30,000 dilution) was immediately added to each well except for non-specific binding wells, followed by 25 μl anticorticosterone antibody (1: 100,000 dilution), and incubated at RT for 2 h. Plates were washed four times (1:20 dilution, 20X Wash Buffer Cat. No. X007; Arbor Assays, Ann Arbor, MI) and 100 μl of TMB substrate solution was added, followed by incubation for 45-60 min at RT without shaking. The absorbance was measured at 405 nm by a microplate reader (TECAN, Sunrise microplate reader, Salzburg, Austria). Assay sensitivity (based on 90% binding) was 0.14 ng/ml. Samples were analyzed in duplicate; intra- and inter-assay CVs were <10% and <15%, respectively.

Concentrations of serum testosterone were quantified by a double-antibody EIA utilizing a polyclonal anti-rabbit testosterone antibody (R156/7; 1:100 dilution) validated for Asian elephants [31]. Samples or testosterone standards (50 μl) were added to appropriate wells. Testosterone-horseradish peroxidase (HRP) (25 μl; 1:10,000 dilution) was immediately added to each well except for non-specific binding wells, followed by 25 μl anti-testosterone antibody (1:110,000 dilution), and incubated at RT for 1 h. Plates were washed four times (1:20 dilution, 20X Wash Buffer Cat. No. X007; Arbor Assays) and 100 μl of TMB substrate solution was added, followed by incubation for 45-60 min at RT without shaking. Absorbance was measured at 405 nm. Assay sensitivity (based on 90% binding) was 0.08 ng/ml. Samples were analyzed in duplicate; intra- and inter-assay CVs were <10% and <15%, respectively.

#### Statistical analysis

Descriptive data were reported as the mean ± standard error of the mean (SEM) and camp management variables were presented as a range or frequency, depending on the type of data. Statistical analyses were performed using R version 3.4.0 [32]. Repeated measures data were analyzed using Generalized Estimating Equations (GEE) to determine: 1) the effects of BCS, FGM and testosterone on metabolic and lipid panel results; 2) seasonal and climate factor effects on metabolic and lipid function; and 3) relationships among metabolic and lipid panel measures. Differences in mean metabolic and lipid profiles between age groups and seasons were further analyzed by Tukey’s post-hoc tests after GEE analyses. Correlations between individual FGM and metabolic hormones or lipid measures in each elephant were analyzed using Pearson’s tests for aggregated data. Mean monthly FGM were compared using GEE followed by a Tukey’s test. Differences in mean FGM, metabolic, lipid profiles and work type between High and Low tourist seasons were analyzed by Tukey’s post-hoc tests. The significance level was set at α=0.05.

## Results

Descriptive BCS, testosterone, FGM, metabolic marker, and lipid profile measures are presented in Table 2, highlighting the variability in mean and range values across individuals. Relationships between BCS, FGM and testosterone on metabolic markers and lipid profiles are presented in Table 3. There were significant positive associations between BCS and TG. FGM concentrations also were positively related to TG, HDL and glucose. Testosterone concentrations were significantly positively associated with HDL, while both BCS and testosterone were negatively correlated to the G:I (Table 3).

**Table 2.** Descriptive statistics. Mean (±SEM) and range values for body condition score (BCS), fecal glucocorticoid metabolite (FGM) and serum testosterone concentrations, lipid panel measures and metabolic factors in captive bull Asian elephants (n = 13) in Thailand tourist camps.

| Parameters | Mean | Min - Max | Mean range |
| --- | --- | --- | --- |
| BCS | 2.80 ± 0.04 | 2.00 - 4.50 | 2.38 - 3.70 |
| FGM (ng/g) | 61.60 ± 2.19 | 1.37 - 330.00 | 37.40 - 98.30 |
| Testosterone (ng/mL) | 2.10 ± 0.18 | 0.03 - 29.39 | 0.51-5.11 |
| TC (mg/dL) | 39.90 ± 0.54 | 19.00 - 101.00 | 28.80 - 46.30 |
| TG (mg/dL) | 26.20 ± 1.11 | 6.00 - 145.00 | 18.20 - 36.70 |
| HDL (mg/dL) | 12.00 ± 0.22 | 2.00 - 29.00 | 7.64 - 17.50 |
| LDL (mg/dL) | 29.20 ± 0.45 | 11.00 - 98.00 | 20.70 - 35.90 |
| Glucose (mg/dL) | 86.10 ± 1.27 | 37.00 - 162.00 | 69.60 - 109.00 |
| Fructosamine (mM) | 0.58 ± 0.01 | 0.38 - 0.82 | 0.54 - 0.62 |
| Insulin (ng/mL) | 0.50 ± 0.05 | 0.03 - 6.82 | 0.05 - 1.59 |
| G:I | 681.00 ± 34.50 | 19.50 - 778.61 | 82.50 - 701.00 |

220 TC = total cholesterol; TG = triglycerides; HDL = high density lipoproteins; LDL = low density lipoproteins; G:I = glucose to insulin ratio.

**Table 3.** General Estimation Equation analyses. Relationships between health factors and body condition and adrenal steroid activity in captive bull Asian elephants (n = 13) in Thailand tourist camps.

| Parameters | BCS |  |  | FGM |  |  | Testosterone |  |  |
| --- | --- | --- | --- | --- | --- | --- | --- | --- | --- |
|  | Intercept | Beta | P value | Intercept | beta | P value | Intercept | beta | P value |
| TC (mg/dL) | 33.47 | 1.610 | 0.270 | 38.88 | 0.013 | 0.110 | 39.66 | 0.146 | 0.350 |
| TG (mg/dL) | 1.87 | 9.280 | 0.009 | 29.09 | 0.101 | 0.009 | 26.93 | -0.155 | 0.570 |
| HDL (mg/dL) | 9.97 | 0.863 | 0.379 | 9.97 | 0.033 | <0.001 | 11.21 | 0.311 | 0.002 |
| LDL (mg/dL) | 24.63 | 0.961 | 0.500 | 29.68 | -0.011 | 0.750 | 29.38 | 0.021 | 0.870 |
| Glucose (mg/dL) | 13.36 | 8.040 | 0.090 | 79.01 | 0.114 | 0.005 | 84.23 | 0.517 | 0.360 |
| Fructosamine (mM) | 0.60 | -0.008 | 0.390 | 0.58 | -2.69-e5 | 0.680 | 0.577 | 0.001 | 0.270 |
| Insulin (ng/mL) | 0.01 | 0.314 | 0.300 | 0.28 | 0.004 | 0.200 | 0.37 | 0.024 | 0.320 |
| G:I | 776.00 | -189.000 | 0.015 | 348.33 | -1.177 | 0.130 | 307.42 | -13.090 | 0.016 |

BCS = body condition score; FGM = fecal glucocorticoid metabolites; TC = total cholesterol; TG = triglycerides; HDL = high density lipoproteins; LDL = low density lipoproteins; G:I = glucose to insulin ratio.

Differences in FGM, testosterone, metabolic marker and lipid profile measures related to age are shown in Table 4. Because of limited numbers, elephants were grouped into two age classes: young (≤30 years) and old (>30 years) for further analysis. Higher levels of TC, TG, LDL, glucose, fructosamine and insulin were found in younger elephants, especially insulin levels, which were three times higher than in older elephants. The G:I was lower in young compared to older elephants. By contrast, testosterone and HDL concentrations were similar across the age groups (Table 4).

**Table 4.** Effect of age on physiological parameters in bull elephants. Mean (± SEM) and range values depicting the effect of age on adrenal and testicular steroid hormones, and health factors in captive bull Asian elephants (n = 13) in Thailand tourist camps.

| Age | FGM<br>(ng/g) | Testosterone<br>(ng/mL) | TC<br>(mg/dL) | TG<br>(mg/dL) | HDL<br>(mg/dL) | LDL<br>(mg/dL) | Glucose<br>(mg/dL) | Fructosamine<br>(mM) | Insulin<br>(ng/mL) | G:I |
| --- | --- | --- | --- | --- | --- | --- | --- | --- | --- | --- |
| Young ($\leq 30$ years)<br>(n = 5) | 60.2±3.44<sup>a</sup><br>14.10-177.00 | 2.18±0.24<sup>a</sup><br>0.05-11.40 | 42.10±0.84<sup>b</sup><br>19.00-101.00 | 30.80±1.84<sup>b</sup><br>6.00-96.00 | 12.40±0.36<sup>a</sup><br>2.00-23.00 | 31.30±0.87<sup>b</sup><br>17.00-98.00 | 98.70±2.14<sup>b</sup><br>47.00-162.00 | 0.58±0.005<sup>b</sup><br>0.38-0.78 | 0.84±0.15<sup>b</sup><br>0.04-6.82 | 204.00±23.30<sup>a</sup><br>19.50-748.00 |
| Old ($> 30$ years)<br>(n = 8) | 70.70±3.12<sup>b</sup><br>14.60-330.00 | 2.05±0.29<sup>a</sup><br>0.03-29.40 | 38.4±0.73<sup>a</sup><br>22.00-101.00 | 23.00±1.44<sup>a</sup><br>7.00-145.00 | 11.80±0.30<sup>a</sup><br>6.00-29.00 | 27.80±0.55<sup>a</sup><br>11.00-55.00 | 77.60±1.36<sup>a</sup><br>37.00-158.00 | 0.57±0.003<sup>a</sup><br>0.49-0.82 | 0.27±0.05<sup>a</sup><br>0.03-2.82 | 336.00±33.20<sup>b</sup><br>51.10-779.00 |

<sup>a,b</sup>Column values are significantly different (p<0.05).
FGM = fecal glucocorticoid metabolites; TC = total cholesterol; TG = triglycerides; HDL = high density lipoproteins; LDL = low density lipoproteins; G:I = glucose to insulin ratio.

Seasonal effects on measured parameters are summarized in Table 5. All but BCS and TG were significantly affected by season. FGM was highest in elephants during the winter months, as was insulin. By contrast, the G:I ratio was lowest during the winter season compared to rainy and summer season months. Glucose also was higher in the winter and summer compared to the rainy season. Serum testosterone was highest in the winter, lowest in the summer, and intermediate in the rainy season. Relationships between environmental factors and BCS, FGM and testosterone concentrations are presented in Table 6, with significant correlations noted between BCS and temperature and humidity. There were no significant negative effects of monthly temperature, rainfall, humidity and THI on FGM or testosterone measures, although an effect of humidity on testosterone approached significance (p = 0.052).

**Table 5.** Seasonal effects on physiological parameters in bull elephants. Mean (±SEM) and range values in body condition scores (BCS), fecal glucocorticoid metabolite (FGM) and serum testosterone concentrations, lipid panel measures and metabolic factors across the summer, rainy and winter seasons in captive bull Asian elephants (n = 13) in Thailand tourist camps.

| Parameters | Summer | Rainy | Winter |
| --- | --- | --- | --- |
| BCS | 2.62 ± 0.06<sup>a</sup><br>(2.00 - 3.00) | 2.75 ± 0.05<sup>a</sup><br>(2.00 - 4.00) | 2.94 ± 0.07<sup>a</sup><br>(2.00 - 4.00) |
| FGM (ng/g) | 54.70 ± 3.64<sup>a</sup><br>(14.90 - 179.00) | 59.5 ± 2.81<sup>a</sup><br>(14.10 - 197.00) | 72.80 ± 4.55<sup>b</sup><br>(14.60 - 166.00) |
| Testosterone (ng/mL) | 1.57 ± 0.25<sup>a</sup><br>(0.07 - 11.40) | 1.96 ± 0.03<sup>ab</sup><br>(0.03 - 29.40) | 2.84 ± 0.45<sup>b</sup><br>(0.18 - 24.00) |
| TG (mg/dL) | 24.40 ± 2.43<sup>a</sup><br>(6.00 - 97.00) | 25.70 ± 1.45<sup>a</sup><br>(9.00 - 90.00) | 28.50 ± 2.41<sup>a</sup><br>(8.00 - 145.00) |
| HDL (mg/dL) | 13.70 ± 0.54<sup>b</sup><br>(6.00 - 29.00) | 11.10 ± 0.27<sup>a</sup><br>(5.00 - 19.00) | 11.90 ± 0.13<sup>a</sup><br>(2.00 - 21.00) |
| LDL (mg/dL) | 27.50 ± 0.72<sup>a</sup><br>(11.00 - 45.00) | 28.80 ± 0.68<sup>ab</sup><br>(16.00 - 59.00) | 31.20 ± 1.09<sup>b</sup><br>(15.00 - 98.00) |
| Glucose (mg/dL) | 88.40 ± 1.80<sup>b</sup><br>(63.00 - 153.00) | 81.00 ± 1.86<sup>a</sup><br>(37.00 - 143.00) | 91.50 ± 3.06<sup>b</sup><br>(47.00 - 162.00) |
| Fructosamine (mM) | 0.61 ± 0.006<sup>c</sup><br>(0.48 - 0.78) | 0.56 ± 0.004<sup>a</sup><br>(0.38 - 0.82) | 0.58 ± 0.004<sup>b</sup><br>(0.49 - 0.68) |
| Insulin (ng/mL) | 0.37 ± 0.08<sup>a</sup><br>(0.04 - 2.89) | 0.33 ± 0.06<sup>a</sup><br>(0.03 - 2.47) | 0.89 ± 0.21<sup>b</sup><br>(0.03 - 6.82) |
| G:I | 290.00 ± 41.60<sup>ab</sup><br>(34.20 - 779.00) | 320.00 ± 33.80<sup>b</sup><br>(40.80 - 702.00) | 186.00 ± 30.80<sup>a</sup><br>(19.50 - 648.00) |

<sup>a,b,c</sup>Row values are significantly different (p<0.05).
TC = total cholesterol; TG = triglycerides; HDL = high density lipoproteins; LDL = low density lipoproteins; G:I = glucose to insulin ratio
Summer: 16 February–15 May, rainy: 16 May–15 October, winter: 16 October–15 February

**Table 6.** General Estimation Equation analysis of seasonal relationships. Relationships between body condition scores (BCS), fecal glucocorticoid metabolite (FGM) and serum testosterone concentrations, and environmental factors in captive bull Asian elephants (n = 13) in Thailand tourist camps.

| Parameters | BCS |  |  | FGM |  |  | Testosterone |  |  |
| --- | --- | --- | --- | --- | --- | --- | --- | --- | --- |
|  | Intercept | Beta | P value | Intercept | Beta | P value | Intercept | Beta | P value |
| Temperature (°C) | 4.410 | -0.057 | 0.048 | 114.300 | -1.810 | 0.093 | 3.895 | -0.065 | 0.248 |
| Rainfall (mm) | 2.832 | 0.007 | 0.900 | 65.714 | -0.450 | 0.620 | 2.098 | 0.001 | 0.980 |
| Humidity (%) | 1.761 | 0.007 | 0.030 | 47.205 | 0.240 | 0.175 | 0.197 | 0.027 | 0.052 |
| THI | 5.144 | -0.030 | 0.175 | 165.570 | 0.864 | 0.132 | 2.790 | -0.009 | 0.840 |

THI = temperature-humidity index

There were significant differences across camps in adrenal activity, metabolic marker and lipid profiles, with FGM, testosterone, BCS, TC, TG, HDL, insulin and glucose being among the highest, and G:I being the lowest in Camp A, the facility with least amount work activities for elephants and the one where tourists fed the highest amounts of treats (e.g., bananas, sugar cane) (Table 7). During the High tourist season, elephants exhibited higher FGM and insulin concentrations than during the Low season (Table 8). In particular, insulin levels were three times higher during the High compared to the Low season, whereas glucose concentrations were unchanged, resulting in a G:I that was 40% lower during the High season.

**Table 7.** Camp differences in body condition, adrenal activity and health markers in elephants. Mean (± SEM) and range values (min-max) for body condition score (BCS), fecal glucocorticoid metabolite (FGM) and serum testosterone concentrations, lipid panel measures and metabolic factors in captive bull Asian in captive bull Asian elephants (n = 13) in three Thailand tourist camps.

| Factor | Camp A | Camp B | Camp E |
| --- | --- | --- | --- |
| BCS | 3.27 ± 0.12<sup>b</sup><br>(2.00 - 4.50) | 2.52 ± 0.10<sup>a</sup><br>(2.00 - 3.00) | 2.62 ± 0.14<sup>a</sup><br>(2.00 - 4.00) |
| FGM (ng/g) | 82.40 ± 3.90<sup>c</sup><br>(21.90 - 179.00) | 58.80 ± 3.40<sup>b</sup><br>(14.90 - 197.00) | 46.50 ± 2.43<sup>a</sup><br>(14.10 - 161.00) |
| Testosterone (ng/mL) | 3.06 ± 0.33<sup>c</sup><br>(0.05 - 16.83) | 2.34 ± 0.53<sup>ab</sup><br>(0.07 - 29.39) | 1.13 ± 0.11<sup>a</sup><br>(0.03 - 6.38) |
| TC (mg/dL) | 43.40 ± 0.75<sup>c</sup><br>(19.00 - 67.00) | 36.20 ± 0.94<sup>a</sup><br>(22.00 - 77.00) | 39.70 ± 1.03<sup>b</sup><br>(27.00 - 101.00) |
| TG (mg/dL) | 31.30 ± 1.96<sup>b</sup><br>(6.00 - 96.00) | 21.90 ± 2.11<sup>a</sup><br>(7.00 - 145.00) | 25.10 ± 1.84<sup>ab</sup><br>(8.00 - 97.00) |
| HDL (mg/dL) | 15.10 ± 0.35<sup>c</sup><br>(7.00 - 23.00) | 11.00 ± 0.43<sup>b</sup><br>(6.00 - 29.00) | 10.20 ± 0.23<sup>a</sup><br>(2.00 - 18.00) |
| LDL (mg/dL) | 29.80 ± 0.65<sup>b</sup><br>(17.00 - 53.00) | 26.90 ± 0.95<sup>a</sup><br>(11.00 - 55.00) | 30.50 ± 0.87<sup>b</sup><br>(20.00 - 98.00) |
| Glucose (mg/dL) | 101.20 ± 2.44<sup>c</sup><br>(47.00 - 162.00) | 73.90 ± 1.54<sup>a</sup><br>(45.00 - 107.00) | 82.00 ± 1.81<sup>b</sup><br>(37.00 - 158.00) |
| Fructosamine (mM) | 0.57 ± 0.005<sup>a</sup><br>(0.41 - 0.71) | 0.57 ± 0.004<sup>a</sup><br>(0.49 - 0.69) | 0.59 ± 0.006<sup>b</sup><br>(0.38 - 0.82) |
| Insulin (ng/mL) | 1.01 ± 0.18<sup>b</sup><br>(0.04 - 6.82) | 0.20 ± 0.05<sup>a</sup><br>(0.03 - 1.59) | 0.20 ± 0.09<sup>a</sup><br>(0.04 - 1.53) |
| G:I | 179.00 ± 26.30<sup>a</sup><br>(19.50 - 748.00) | 305.00 ± 52.30<sup>ab</sup><br>(65.50 - 701.00) | 349.00 ± 33.00<sup>b</sup><br>(69.00 - 779.00) |

<sup>a,b,c</sup>Row values are significantly different (p<0.05).
TC = total cholesterol; TG = triglycerides; HDL = high density lipoproteins; LDL = low density lipoproteins; G:I = glucose to insulin ratio

**Table 8.** Tourist season effects on physiological parameters in bull elephants. Mean (± SEM) and range values for captive bull elephants in lipid profiles and metabolic factors between High and Low tourist seasons in captive bull Asian elephants (n = 13) in Thailand tourist camps.

| Factor | High season | Low season |
| --- | --- | --- |
| BCS | 2.98 ± 0.07<sup>a</sup><br>(2.00 - 4.50) | 2.70 ± 0.04<sup>a</sup><br>(2.00 - 4.00) |
| FGM (ng/g) | 73.50 ± 5.07<sup>b</sup><br>(14.60 - 330.00) | 60.40 ± 2.33<sup>a</sup><br>(14.10 - 197.00) |
| TC (mg/dL) | 41.80 ± 1.14<sup>a</sup><br>(22.00 - 109.00) | 39.10 ± 0.59<sup>a</sup><br>(10.00 - 85.00) |
| TG (mg/dL) | 26.30 ± 2.18<sup>a</sup><br>(7.00 - 145.00) | 26.10 ± 1.27<sup>a</sup><br>(6.00 - 97.00) |
| HDL (mg/dL) | 12.20 ± 0.38<sup>a</sup><br>(2.00 - 22.00) | 12.00 ± 0.27<sup>a</sup><br>(5.00 - 29.00) |
| LDL (mg/dL) | 30.30 ± 1.01<sup>a</sup><br>(11.00 - 98.00) | 28.70 ± 0.46<sup>a</sup><br>(12.00 - 59.00) |
| Glucose (mg/dL) | 89.30 ± 2.59<sup>a</sup><br>(47.00 - 162.00) | 84.70 ± 1.41<sup>a</sup><br>(37.00 - 153.00) |
| Fructosamine (mM) | 0.58 ± 0.01<sup>a</sup><br>(0.48 - 0.78) | 0.58 ± 0.01<sup>a</sup><br>(0.38 - 0.82) |
| Insulin (ng/mL) | 0.90 ± 0.13<sup>b</sup><br>(0.03 - 6.82) | 0.35 ± 0.03<sup>a</sup><br>(0.03 - 2.89) |
| G:I | 186.00 ± 15.70<sup>a</sup><br>(19.50 - 648.00) | 307.00 ± 13.80<sup>b</sup><br>(34.20 - 779.00) |

<sup>a,b,c</sup>Row values are significantly different (p<0.05).
BCS = body condition score; FGM = fecal glucocorticoid metabolites; TC = total cholesterol; TG = triglycerides; HDL = high density lipoproteins; LDL = low density lipoproteins; G:I = glucose to insulin ratio.
270 High: November – February, Low: March – October

Several correlations were noted amongst the metabolic and lipid factors as shown in Table 9. Strong positive associations (p<0.001) were found between TC and HDL, LDL and HDL and glucose, and glucose and insulin. By contrast, negative relationships (p<0.001) were found between the G:I and HDL and glucose, and between insulin and G:I. In separate Pearson’s correlation analyses of individual means (n = 13), FGM levels were similarly correlated to HDL, glucose and insulin, and negatively correlated to G:I (p<0.05) (Fig. 1).

**Table 9.**
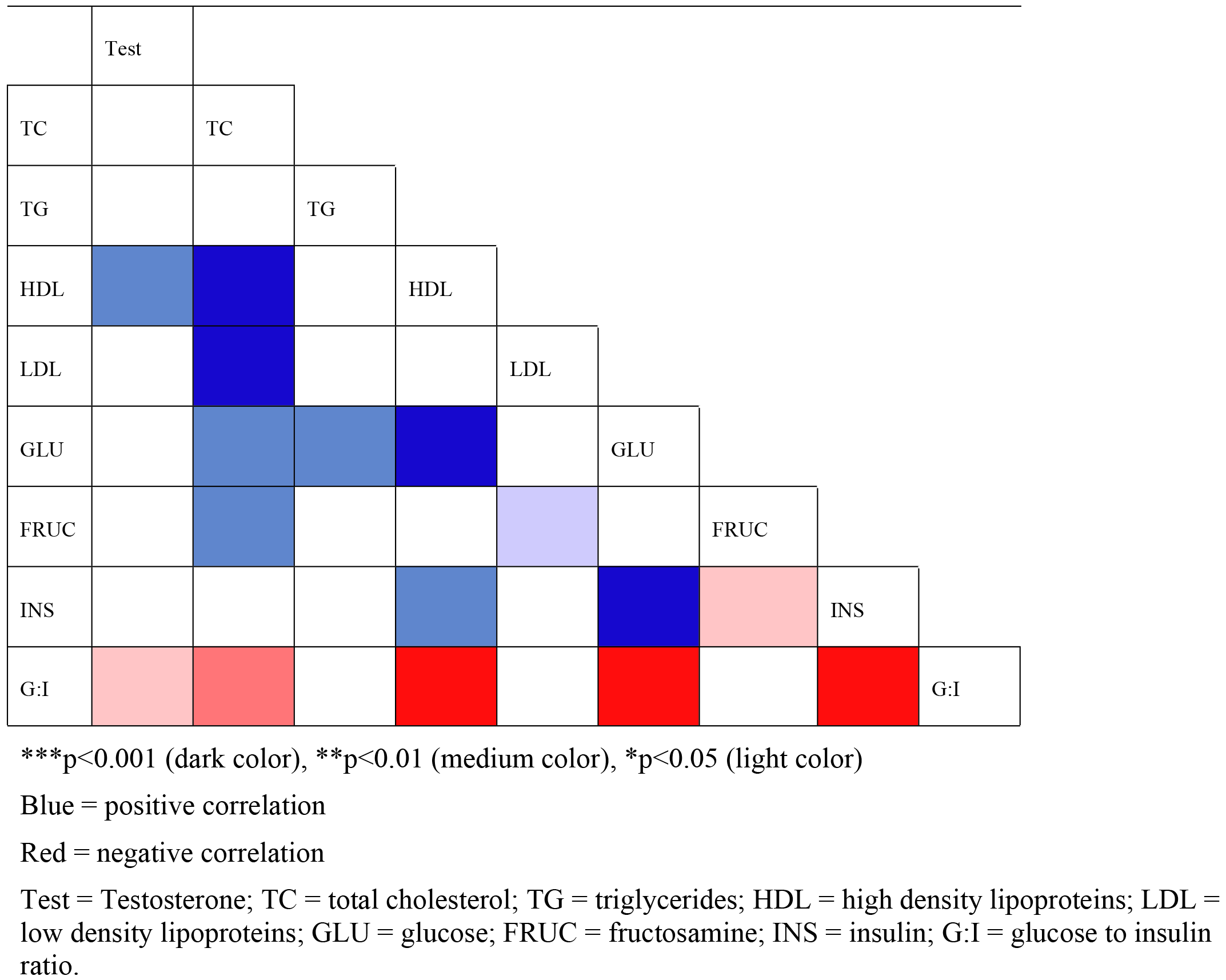
Relationships among physiological factors in bull elephants. Correlation matrix presenting relationships between testosterone and lipid panel measures and metabolic factors in captive bull Asian elephants (n = 13) in Thailand tourist camps.

**Figure 1.**
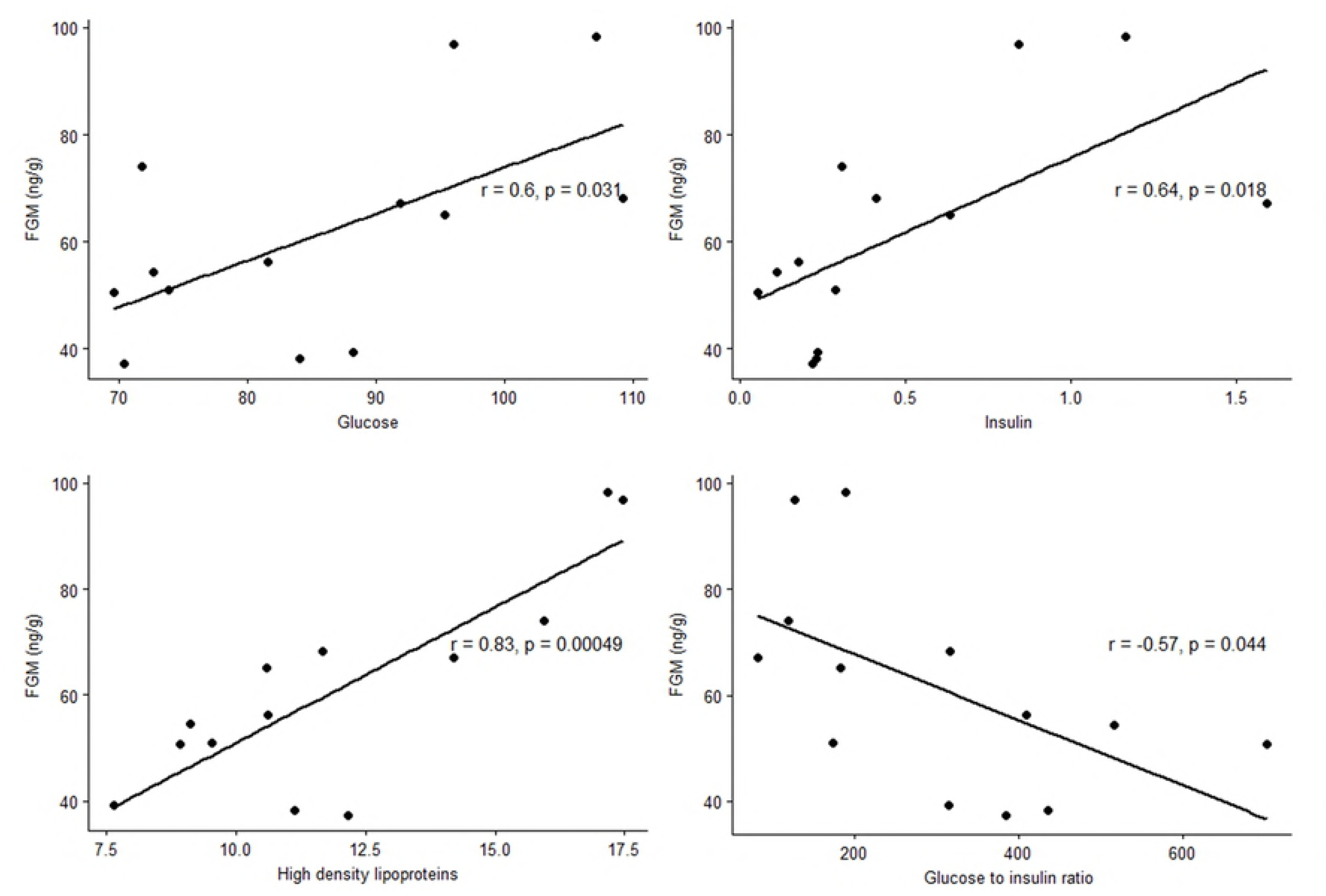
Significant relationships between adrenal steroids and lipid and metabolic factors. Pearson’s correlations between FGM concentrations and lipid panel measures and metabolic factors in captive bull Asian elephants (n = 13) in Thailand tourist camps.

## Discussion

This was the first study to assess lipid profiles, and evaluate relationships between BCS, FGMs and metabolic factors in male Asian elephants in Thailand. The mean BCS was 2.8, which is within a normal/ideal distribution of body fat for elephants [33]. In general, the working bulls in our study had better body condition than western zoo bull elephants; 69% of Thai bulls had a BCS = 3, compared to only 33% in the U.S. Furthermore, 48% of bulls in the U.S. had a BCS = 4, whereas no bulls in Thailand had an average BCS of 4 or 5 over the study period, although a few scored up to 4.5 in individual months. This could be due to higher amounts of exercise, with tourist elephants engaged in many activities, including trekking, bathing, shows or walking with tourists [34], so inactivity is less of a concern. This result is consistent with a study of female Asian elephants in Thailand where most (~60%) had a BCS = 3 [11], although compared to males, females had on average higher BCSs than males (3.50 + 0.02 vs 2.80 + 0.04; GEE, p<0.001). In the U.S., elephants that walked more than 14 hours/week had a decreased risk of BCS = 4 or 5 [35]. Thus, increased exercise associated with tourist activities likely helps elephants maintain better body condition. From a previous study, female elephants that worked more hours per day in the form of saddle or bareback riding were found to have better BCSs and metabolic health [11].

Insulin plays a central role in the regulation of blood glucose and energy homeostasis; however, high levels are associated with hypertension, obesity, dyslipidemia, and glucose intolerance in humans [36, 37]. Glucose and insulin are generally measured after a patient has fasted, but that was not possible in our elephants because they had access to forage overnight. Instead, we evaluated the G:I [38], which has been used to detect of insulin sensitivity in women [10]. One popular activity for tourists is feeding, particularly bananas and sugar cane, which possess high concentrations of sucrose and other soluble sugars that could contribute to weight problems. Blood glucose values agreed with the previous study of female elephants (86.10 + 1.27 vs 88.90 + 0.75 mg/dL) that found elephants with a BCS of 4.5 - 5.0 had higher glucose and insulin levels, and a lower G:I [11]. Compared to females at the same camps, bulls exhibited a higher, and thus potentially healthier, G:I (681.00 + 34.50 vs 196.00 + 6.72; GEE, p<0.001). Last, the G:I of bulls in Thailand was higher than that of bulls in the U.S. (253 ± 18) [4], commenserate with the higher BCSs observed in the latter population. These results agree with insulin measures, which were lower in male than female elephants (0.50 + 0.05 vs 0.75 + 0.03 ng/mL; GEE, p=0.003), respectively [11]. Thus, overall, it appears the bull elephants in our study population are healthier than their female cohorts, both physically and physiologically, perhaps because feeding of bulls is more limited. They also differ somewhat in metabolic profiles compared to bulls in the U.S., which deserves further study.

Serum TG concentrations were within the range reported for female Asian elephants [2, 11] and correlated positively with BCS, consistent with other findings [11, 35]. From work in other species, serum TGs are useful indicators of overall adiposity [39–42]. In humans, a number of metabolic changes are associated with obesity, high BMI or poor eating habits, including elevated TC, TG and LDL levels. Similarly, in dogs, increased plasma TC and TG concentrations are observed in association with obesity [3], and in horses, TG concentration is correlated with body condition [43, 44]. Hyperlipidemia could pose problems for elephants, although incidence of cardiovascular disease appears to be relatively low [45], and none of the bulls in our study were obese (BCS = 5).

There were positive associations between FGM and TG, HDL and glucose, indicating relationships between adrenal activity and metabolic and lipid function, as has been demonstrated in other species [46, 47], including female elephants [11]. Elevated and sustained cortisol secretion during chronic stress can lead to central obesity, hypertension, glucose intolerance, and dyslipidemia in humans [48]. Studies to understand how management factors affect stress responses in captive animals are key to improving welfare, and are beginning to be applied to elephants in westerns zoos [49, 50], and to working tourist elephants in Asia [11]. In exploring age effects, older bull elephants (>30 years) exhibited comparatively lower metabolic and lipid function than those under 30 years, in agreement with studies in other species that show basal metabolic rate decreases almost linearly with age [51–53]. By contrast, in humans, diabetes is a progressive disease, and glucose levels tend to increase with age. The incidence of type-2 diabetes increases in older individuals primarily due to age-related declines in beta cell function and impaired insulin secretion, rather than to insulin resistance [54]. The aging process can also be associated with pancreatic islet cell dysfunction as a contributing factor to abnormal glucose metabolism [55]. We are aware of only one study of diabetes in an older Asian elephant bull [56], so this condition may not be common. Nevertheless, these results highlight a number of physiological changes associated with age in this species.

There was a significant effect of season on FGM, with higher concentrations during the winter when temperatures and rainfall are lower. Salivary cortisol and FGM in female Asian elephants in Spain and Thailand, respectively, were highest during the winter [11, 57]. In goral, another indigenous ungulate species in Thailand, FGMs also were higher in winter [58, 59]. The need for more energy to maintain optimum body temperature and ensure survival in cooler temperatures could be related to this finding. Elevated circulating GC levels as a response to cold stress have been documented in red deer (*Cervus elephas*) [60] and in farm animals [61]. However, it should be noted that the winter months also correspond to the High tourist season, so more work is needed to identify the primary drivers of adrenal activity in working elephants.

Significant differences were identified across camps in stress hormone levels, metabolic status and lipid profiles, which may be related to management and tourist activities. Elephants at Camp A exhibited overall lipid and metabolic results that were higher on average than the other camps. BCS at Camp A also was the highest, while G:I was the lowest, indicative of metabolic derangements [4, 11]. These effects appear to be related, in part, to the feeding of greater amounts of supplementary foods (30 vs 5 - 10 kg/day) and lower levels of activity (1.1 vs 3 - 4 hours/day). Female elephants at Camp A also exhibited higher BCS, FGM concentrations, and metabolic and lipid measures, suggesting management changes may be needed to provide elephants with better diets or increased exercise opportunities. Some of these camp effects may be related to tourists, given that there was a significant tourist season effect on health status, with levels of FGMs and several metabolic markers being higher during the High tourist season. Higher numbers of tourists likely are associated with increases in amounts of food treats offered to elephants, given that feeding is one of the most popular activities. The food given to elephants in this study consisted of items with a high sugar content and glycemic index, including bananas, sugarcane, watermelon, and pumpkin. High glycemic index foods induce an exaggerated insulin response, which can increase body fat and weight, and lead to insulin resistance, and eventual exhaustion of endocrine pancreatic function and insulin release [62, 63]. There is growing recognition and concern that obesity and metabolic conditions are negatively impacting the health of many species, including humans, companion and domestic animals. A similar health concern exists for zoo-held species, including elephants, that often are fed diets high in calories and given inadequate exercise [2, 4, 33]. However, for the bulls in this study, body condition, adrenal activity, metabolic and lipid status all appeared to indicate a comparatively normal health status.

By contrast, compared to females at the same camps [24], overall FGM concentrations in bulls were higher overall (Camp A, 82.40 ± 3.90 vs 60.40 ± 2.43; Camp B, 58.80 ± 3.40 vs 49.60 ± 2.40; Camp E, 46.50 ± 2.43 vs 39.60 ± 2.06 ng/g, GEE, p<0.001), respectively. One explanation could be that bulls are controlled more vigilantly (i.e., using an ankus) around tourists, and are often kept on shorter chains because of their more aggressive nature [64], which could lead to higher stress levels. A big challenge in captive elephant management is how to care for bulls in a way that meets their welfare needs; a problem that often is ignored because of limited capacity to safely and humanely manage them.

Testosterone serves a number of biological functions, including development of male reproductive tissues, promoting secondary sexual characteristics, and supporting breeding behaviors. However, low testosterone in men and animals has been associated with a metabolic syndrome characterized by obesity, diabetes, hypertension, and dyslipidemia [65–68], and clinical trials have demonstrated that testosterone replacement therapy moderates problems with insulin resistance and aids in glycemic control [69]. Although none of the bulls in our study were overweight or obese (mean BCS = 4 or 5), there was a positive correlation between testosterone and HDL (good cholesterol), in agreement with previous studies in humans [70, 71], and a negative relationship with G:I, suggesting thinner bulls may experience better health parameters, not unlike our studies in female elephants. We did note a seasonal effect on serum testosterone. Three of the bulls came into musth between October and December, for 2 - 3 months each, which is the general pattern for elephants in Thailand [72, 73]. Although blood samples were not collected during musth due to safety concerns, average testosterone concentrations were still higher in winter as compared to summer and rainy season months, presumably reflecting changes in baseline concentrations. A study of bulls in Thailand by Thongtip et al. [73] also found higher serum testosterone during the winter, despite not collecting samples during musth. Overall, there was no difference in testosterone concentrations in bulls >30 years, which differs from that of Brown et al. [74], who found an overall increase with respect to age (r = 0.69) in Asian bulls in the U.S., at least up to the age of 42 years. That study did include samples from musth bulls, which could account in part for the study differences.

Compared to the females in Norkeaw *et al*. [11], there were similar positive relationships between TC and HDL and LDL, HDL and glucose, glucose and insulin, and negative relationships between HDL and G:I, glucose and G:I, and insulin and G:I. By contrast, males differed from females by exhibiting a negative correlation between TC and G:I, and a lack of positive relationships between glucose and fructosamine, and fructosamine and insulin. In the cat, fructosamine concentrations are tightly correlated with blood glucose levels in obese animals [75], but there was no such relationship in bull elephants, likely because we saw no incidence of obesity in this population.

## Conclusion

Used as outcome variables in regression models, greater BCS and FGM measures were predictors of higher metabolic and lipid levels in male Asian elephants, although not to the extent observed in female elephants in these same camps. There also was a relationship between higher BCSs and adrenal steroid hormone measures in bulls during the winter, when there are more tourists. Thus, elephant health and well-being could be promoted by limiting tourist interactions with individual elephants, especially during the High season. However, it may be significant that overall, bull elephants in Thailand appear to be in better physical and physiological health compared to females at the same camps. This difference could be related to how bulls are managed, keeping them at a lower body condition to lessen musth symptoms and perhaps to fewer feeding opportunities with tourists.

## Acknowledgements

We are grateful to the elephant owners and mahouts for participating in this study and allowing us to work with the elephants. We would like to thank our colleagues, Dr. Muyao Li, Ms. Patcharapa Towiboon, Mr. Pallop Tankaew, Dr. Khajohnpat Boonprasert, Dr. Patiparn Toin, Dr. Tithipong Plangsangmas, Dr. Channarong Saisaard, Dr. Panida Muanghong and Dr. Siripat Khammesi for help with collecting blood samples and for laboratory assistance.

